# Social context modulates idiosyncrasy of behaviour in the gregarious cockroach *Blaberus discoidalis*

**DOI:** 10.1101/028571

**Authors:** James D. Crall, André D. Souffrant, Dominic Akandwanaho, Sawyer D. Hescock, Sarah E. Callan, W. Melissa Coronado, Maude W. Baldwin, Benjamin L. de Bivort

## Abstract

Individuals are different, but they can work together to perform adaptive collective behaviours. Despite emerging evidence that individual variation strongly affects group performance, it is less clear to what extent individual variation is modulated by participation in collective behaviour. We examined light avoidance (negative phototaxis) in the gregarious cockroach *Blaberus discoidalis*, in both solitary and group contexts. Cockroaches in groups exhibit idiosyncratic light-avoidance performance that persists across days, with some individual cockroaches avoiding a light stimulus 75% of the time, and others avoiding the light just above chance (i.e. ~50% of the time). These individual differences are robust to group composition. Surprisingly, these differences do not persist when individuals are tested in isolation, but return when testing is once again done with groups. During the solo testing phase cockroaches exhibited individually consistent light-avoidance tendencies, but these differences were uncorrelated with performance in any group context. Therefore, we have observed not only that individual variation affects group-level performance, but also that whether or not a task is performed collectively can have a significant, predictable effect on how an individual behaves. That individual behavioural variation is modulated by whether a task is performed collectively has major implications for understanding variation in behaviours that are facultatively social, and it is essential that ethologists consider social context when evaluating individual behavioural differences.

**Abbreviations:** DLP: digital light processing; MP: megapixels; ANOVA: analysis of variance, ICC: intraclass correlation coefficient, CI: confidence interval.

---

In animal groups, individuals with different phenotypes can nevertheless coordinate their behaviours to solve problems and increase individual fitness. Group living increases the chance of encountering a mate (Uzsak & Schal, 2013), provides security from predators (Uzsak & Schal, 2013; Treherne & Foster, 1980), and enhances access to other key resources such as food and shelter (Parrish et al., 1999). Group dynamics are important for understanding how animals use collective decision making to solve problems and attain high levels of fitness.

To understand group dynamics, we need to examine the relationship between individual variation and collective behaviour. This relationship is complex, however, and is currently a frontier of research in animal behaviour (Bengston and Jandt 2014, LeBoeuf and Grozinger 2014, Jeanson and Weidenmüller 2014, Jandt et al 2014). It is clear that individual variation (arising through a number of mechanisms, including genetic diversity (Bengston and Jandt 2014), or differences in experience (Ravary et al 2007)) can give rise to variation between groups through a variety of processes, such as founder effects or interactions with conspecifics, etc. (Bengston and Jandt 2014, LeBoeuf and Grozinger 2014). Increasingly, however, there is also evidence that the presence of conspecifics can drive individual behavioural variation (LeBoeuf and Grozinger 2014), for example through social niche differentiation (Bergmüller and Taborsky 2010). Individual variation can thus affect, but also be affected by, group behaviour.

There is strong empirical evidence for individual variation in collectively behaving animals. Schools of fish (Marras & Domenici, 2013), flocks of homing pigeons (Hoffman 1958), and even human groups (Vindenes et al., 2008) are populated by highly varied individuals, which can have important effects on group performance (Brown and Irving 2013). Among invertebrates, castes within eusocial insects are a classical example of behavioural differentiation within a group context (O’Donnell 1998; Winston & Michener, 1977). These differences can emerge even when all individuals are genetically identical (Freund et al., 2013), suggesting that individual variation in behaviour could be an emergent property of group membership. Yet, eusociality is not a prerequisite for behavioural differences between individuals. Indeed, several non-eusocial insects exhibit conspicuous individual differences even when genetically identical (Petrokvsii et al., 2011; Schuett et al., 2011; Kain et al., 2012; Stamps et al., 2013; Buchanan et al., 2015), likely reflecting developmental noise rather than an emergent property.

As an intermediate case between eusocial and solitary lifestyles, gregarious insects represent an interesting case for the consideration of individuality in the group context. Clonal, gregarious aphids exhibit individuality in both escape (Schuett, et al., 2011) and exploratory locomotion behaviours (Petrovskii et al., 2011). Canonge et al. (2009) showed that American cockroaches *(Periplaneta americana*) exhibit individual differences in resting site preferences. Planas-Sitjà et al. (2015) found (in the same species) that behavioural variation between individuals can affect group dynamics and collective shelter-seeking behaviour. However, the interplay between individual variation and collective behaviour in gregarious insects remains a nascent research area.

There is emerging evidence that such individual variation plays an important role in determining collective behaviour (Hui and Pinter-Wollman 2014, Modlmeier et al 2014a) and group success (Pruitt and Reichert 2011, Modlmeier, Liebmann, and Foitzik 2012). Individual variation in social spider groups (*Stegodyphus dumicola*) plays a larger role in determining group success than the size of the group (Keiser and Pruitt, 2014). Hoffman (1958) showed that even in humans, the individual variation within a group significantly contributes towards that group’s success. The effect of individual differences on group behaviour can be distributed evenly across individuals or concentrated in specific members. Key individuals in a group can have a particularly strong influence on the collective behaviour of their group (Modlmeier et al., 2014b).

Despite increasing evidence that individuality plays a large role in determining collective behaviour, we have only recently begun to understand the potential effects of group membership in modulating individual variation. In social spiders, group membership can increase individual behavioural variation (Laskowski and Pruitt 2014, Modlmeier et al. 2014c). In social insects, there has been increasing interest in understanding how feedback between individual behaviour and social context may dynamically produce stable, individually specific behavioural patterns (Bengston and Jandt 2014, Jandt et al. 2014, Jeanson et al. 2014, LeBoeuf et al 2014). In honeybees, for example, colony context has a clear effect on at least some behaviours, with clonal subpopulations of bees exhibiting different behavioural patterns depending on the genetic homogeneity of the entire colony (Hunt et al 2013, Gempe et al 2012). Outside of social insects, there is also evidence that social context can modulate behavioural traits typically associated with “personality” (i.e. risk-taking behaviour, van Oers et al 2005, Schuett et al. 2009, or “boldness”, Keiser et al. 2014). However, the extent to which such group effects are pervasive outside of highly social arthropods is largely unknown.

Our broad goal is to use cockroach light-avoidance behaviour to examine 1) how individual behavioural differences correlate with collective behaviour in a system that allows rapid quantification and robust tracking of individuals across contexts and 2) the effect that group membership has on individual variation. Cockroach light-avoidance is likely a predator-evasion and shelter-seeking response. Performance (defined as the fraction of time spent in the shade) of this behaviour improves with the size of the group, and thus can be considered a collective behaviour (Canonge et al., 2011; Salazar et al., 2013; Sempo et al., 2009). When searching for a suitable shelter, cockroaches are able to use social cues to reach a consensus and aggregate in a single suitable shelter (Sempo et al., 2009). However, the consensus decision is influenced by the individual variation within a group (Sempo et al., 2009). Thus we expect to also find that individual variation in light-avoidance performance contributes to differences at the group level.

Using a new 2D bar-coding system (Crall et al., 2015) we tracked individual cockroaches as they performed a collective light-avoidance behaviour, in a variety of group configurations, to test the following hypotheses. First, we hypothesize that individual animals will display different behaviours with respect to the light stimulus. Specifically, some individuals will be better at avoiding the light than others. We also hypothesize that these differences between individuals emerge from social niche construction occurring after the formation of those experimental groups. We re-assigned individual roaches from their original random groups to groups based on similarity in their individual light-avoidance performance. If social niche construction acts on days-long timescales, individual variation in performance would re-emerge even in groups initially composed of individuals with little variation. These experiments assess the stability of individual differences across changes in group-membership. Next, using solitary light-avoidance assays, we tested the hypothesis that any stable individual differences observed across the first two experiments would persist when animals were assayed individually. Finally, by restoring the animals to experimental groups, we tested the hypothesis that any discrepancy between individual behaviours in the group and solitary contexts could be explained by drift in individual behavioural biases over time.

## Materials and Methods

We developed a system for automatically tracking cockroach position in a circular arena, in which a downward-facing projector delivered a moving light/shade stimulus, and cockroach position was imaged using light invisible to the cockroaches. Cockroaches were permanently tagged with optical codes whose position could be extracted from the frames of a video using pattern recognition software (Crall et al., 2015). Combining these two techniques, we were able to determine a cockroach’s position, speed, and whether it was in the light or the shade. The use of permanent tags enabled us to track the performance of individual cockroaches over a month of successive experiments, even while varying the membership of the groups.

Scripts and processed cockroach position data are available at: *http://lab.debivort.org/social-context-modulates-idiosyncrasy* and Dryad.

### Study organism and animal care

*Blaberus discoidalis* animals were purchased from Backyard Brains and were approximately 8 months old on arrival. 60 males free of conspicuous external damage were taken from a mixed sex population and used as experimental individuals. Cockroaches were housed in opaque black plastic containers with translucent white perforated lids. Houses contained egg-carton cardboard enrichment. Food pellets (Rat and Mouse food, Carolina Biological Supply) and water-soaked paper towels were replaced weekly. Containers were cleaned weekly.

The test cockroaches were tagged with BEEtag codes for automated video tracking (Crall et al., 2015). Tags were printed on waterproof paper and were ~ 8 × 8 mm in size. Each cockroach was anaesthetized using CO_2_. While anaesthetized, the pronotum of the cockroach was abraded with fine grit sandpaper and the BEEtags were attached to the pronotum using cyanoacrylate glue. Cockroaches were given a minimum of two days to recover after anaesthetization before the start of experimental trials. During this time 2 out of ~80 tagged individuals shed their tags, and were not retained for experiments.

### Experimental set-up and stimulus

A circular arena with walls made from high-density polyethylene was constructed by cutting the top and bottom off a 5-gallon liquid waste container. The circular arena was 28.2 cm in diameter and ~30 cm tall. A mounting base for the arena was constructed with black 5.6 mm acrylic. The arena walls could be slotted into a ~5 mm wide circular groove cut into this base, holding the walls in place. For trials, the base was covered by a sheet of Absorbent Lab Paper (VWR-51138-500), which was changed between trials, to minimize odorant contamination. An Optoma S316 DLP (Digital Light Processing) projector and 5MP monochromatic digital camera with a global shutter (Blackfly model, Point Grey) were mounted on an aluminum extrusion rig above the arena. Recordings were collected at 7 frames per second, with an exposure time of 8 ms. This exposure time was chosen to minimize motion blur within each frame, as well as to synchronize with the vertical scan of the DLP projector. The projector delivered a computer-controlled stimulus onto the base and interior walls of the arena. The camera was covered by a 590 nm long pass red filter (Thorlabs). The camera recorded a video of the entire base of the arena for the duration of each trial.

For tagging control experiments (Fig S1), the projected stimulus was magenta on the top half and red on the bottom half. Control experiments were conducted with individual cockroaches (n=21 untagged, 19 tagged) and in groups of three randomly selected individuals (*n*=6 untagged groups, 5 tagged groups). For all other experiments, the projected stimulus was alternating red and magenta quadrants (Fig 1). The quadrants rotated at 0.05Hz and randomly reversed rotation direction with probability 0.333 per frame, resulting in an average rotational direction persistence of 1s. The stimulus also included two small black wedges at the center of the red sectors, which allowed us to use machine vision to identify the position of the sectors in the same image we used to track the BEEtags. These colors were chosen because cockroaches do not perceive red light, so the red segments of the stimulus would appear to be dark to them while the magenta stimulus would appear bright (Walther, 1958). Before each trial, experimental animals were transferred to an empty plastic container in darkness. The stimulus was then initiated and the cockroaches were gently poured into the arena (Movie S1). Recordings lasted either 10 or 5 minutes.

**Figure 1.**
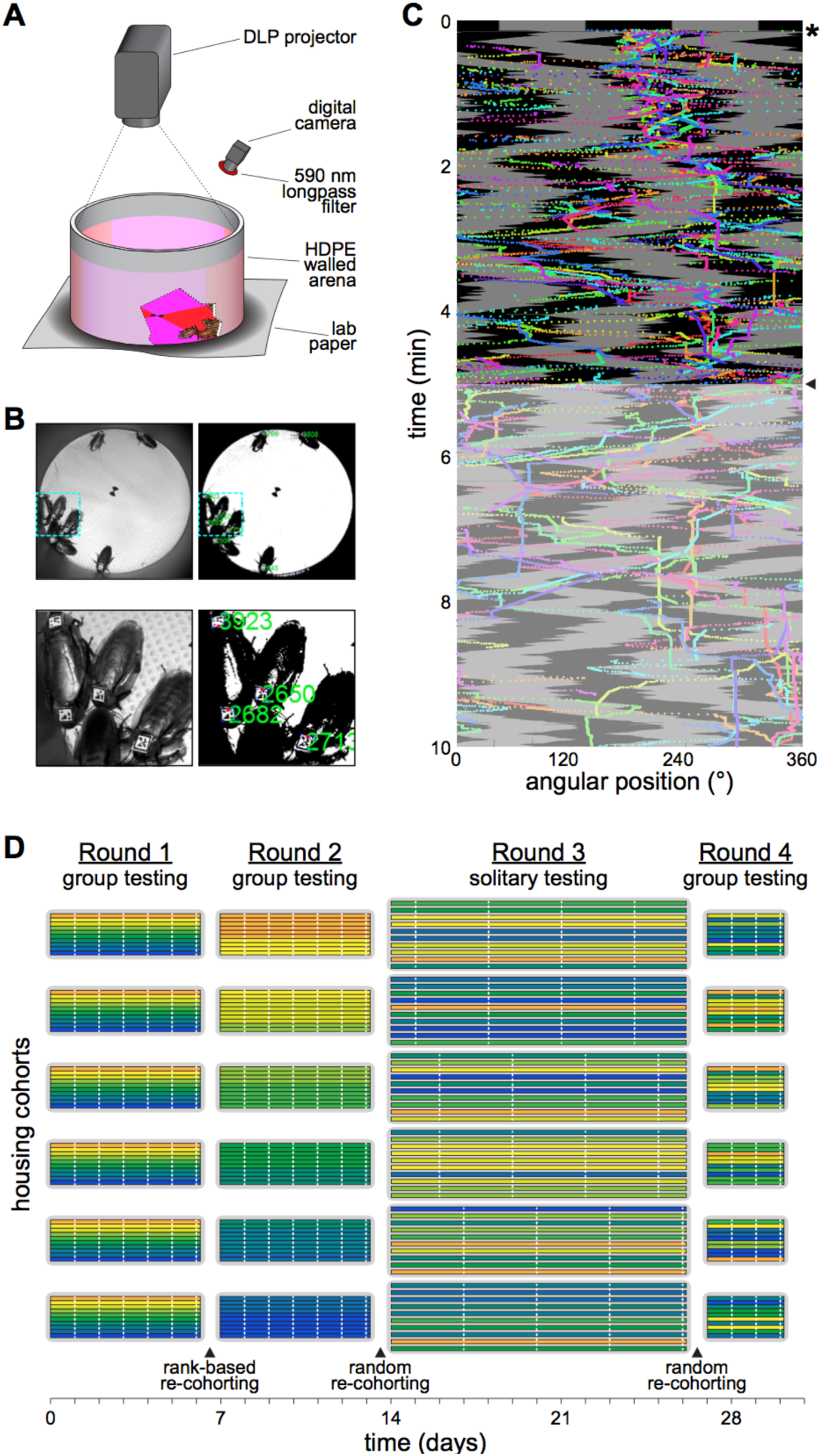
An automated tracking system monitors individual cockroach behaviour during group phototaxis. **A)** Diagram of experimental setup. Circular arena resting on absorbent lab paper directly under a projector that projects the moving stimulus onto the arena. A digital camera is positioned to capture the entire arena, this camera is filtered with a 590 nm long pass filter to allow digital tracking through both light conditions. **B)** Upper row: A still frame of the tracking video during a group trial (left) and the same image showing the number and location of identified tags. Lower row: Cropped images from each of the upper row panels. **C)** Kymograph with time running along the vertical axis, depicting cockroach angular position in the arena as it relates to the angular position of shaded and lit regions over time. The black region corresponds to the red-zone of the arena. Each cockroach has a unique colour trail throughout the timeline of the kymograph. It is important to note that only the top half, or the first five minutes, of the kymograph are used in the analysis. **D)** The cohort composition during each of the four rounds of trials. Round two cohorts are based on tracking performance in round one while the other round’s cohorts were randomly selected. Colors illustrate hypothetical tracking performance.

### Trial structure

We conducted four different experiment phases (Rounds), varying the composition of housing and experimental groups (cohorts).

*Round 1:* In the first round of trials, 60 individuals were randomly placed into 6 cohorts of 10 cockroaches each. To ensure that cohort composition was not influenced by the relative ease of picking up some cockroaches compared to others, we took a population of 80 tagged cockroaches and divided them into 5 groups of equal size. Cockroaches were placed in these temporary groups in the order they were picked up in (the first 16 cockroaches picked up went into group 1, second 16 went into group 2 etc.). From each group we randomly selected two cockroaches to be in each of the experimental cohorts. The cockroaches were allowed to acclimate to their new housing group for two days. Each 10-individual cohort then underwent one experimental trial each day on 6 consecutive days, in which the entire cohort was introduced into the arena for a 10-minute trial (Fig 1D). Here and in all analyses, tracking performance was defined as the percentage of time that each individual spent in the red zones. Only the first 5 minutes was considered because the cockroaches habituated to the stimulus (see below).

*Round 2:* After the last experiment of Round 1, the cockroaches were then placed in new housing and experimental cohorts (“re-cohorted”) based on their ranked individual tracking performance in the first round. Rank 1 individuals were all added to the first new cohort. Rank 2 individuals were randomly split between the first and second Round 2 cohorts, so that the first cohort had 10 members, etc. This procedure was continued to populate all Round 2 cohorts (Fig 1D). The cockroaches were then given 48 hours to acclimate to their new housing groups. Experiments in Round 2 proceeded as in Round 1, with each cohort of 10 individuals being tested six times.

*Round 3:* After the last Round 2 experiment, the cockroaches were re-cohorted randomly into 6 new cohorts with the use of a 6-sided die. The cockroaches were then given 48h to acclimate to their new housing groups. For experiments, each individual was introduced alone into the arena (Movie S2), and its movements recorded for a total of 5 minutes. The stimulus presentation during trials was identical. Each day cockroaches from two cohorts were tested, and this was repeated until each individual was tested 4 times in this Round, which consequentially lasted for 12 days (Fig 1D).

*Round 4:* After the last Round 3 experiment, the cockroaches were re-cohorted randomly into 6 new cohorts with the use of a 6-sided die. The cockroaches were then given 48h to acclimate to their new housing groups. These cohorts underwent group trials similar to the trials described in the first and second rounds of trials. Each cohort of 10 individuals was tested three times each over the course of Round 4.

Thus, in terms of fully independent units (i.e. the sample sizes), Rounds 1 through 4 had, respectively, 6 cohorts of 10 roaches each, 6 cohorts of 10 roaches each, 60 individual roaches and 6 cohorts of 10 roaches each. The number of replicate experiments per Round was, respectively, 6, 6, 4, and 3. During the 31 days of experiments, three cockroaches died, the first between trials 1 and 2 of the first round of experiments. It was replaced with a randomly chosen individual from the pool of tagged cockroaches. The subsequent cockroaches were not replaced, so at any time, up to two experimental and housing cohorts had 9 rather than 10 individuals.

### Automated behavioural analysis

All recordings were saved in raw monochromatic .avi format and processed using custom scripts in MATLAB. For control experiments comparing tagged and untagged cockroaches, movies were imported, and 50 frames evenly spaced throughout the recording were collected. These were combined using a median filter to generate an image of the empty arena for background subtraction. Subtracted images were thresholded and noise was reduced by eroding and dilating above threshold pixels until only cockroaches’ outlines remained. In solitary control trials the center of a convex hull surrounding the cockroach outlines was taken as the animal’s positions. In 3 animal trials, the aggregation index was calculated as the area enclosed by the convex hull surrounding all outlines.

BEEtag positions were extracted from thresholded images using published code (Crall et al., 2015). For each trial, the center of the arena was marked manually upon running the MATLAB script. To determine an optimal image threshold for tag identification, a sample of frames from throughout the recording was chosen and thresholds systematically varied. The threshold identifying the greatest number of tags from those sample frames was used for the whole movie. Based on the indicated center of the arena, the cockroach positions were translated into polar coordinates, and the angular position chosen as the focus for analysis. Sometimes tracking of the position of the red sectors generated errors, e.g., when a cockroach walked over the small black targeting sectors. To address this we used an interpolation script to make a “best guess” estimate of the sector positions for each frame. Individuals were untrackable on some frames due to motion blur, foreshortening of the BEEtags, being obscured by other cockroaches, being flipped upside down, or (rarely) walking through the unilluminated black targeting sectors. We replaced these missing values with values linearly interpolated across the gap of missing values (Movie S3). Average instantaneous speeds for each cockroach were calculated as a proxy for activity. Average velocity within a trial was highly correlated with portion of time spent moving, since cockroaches had a relatively characteristic speed when moving, and we therefore only included average speed in our analyses here.

### Statistics

ANOVAs and regression analyses were calculated in MATLAB or R with built-in functions. For all ANOVAs, individual cockroaches provided the independent grouping variables. Boxes in box plots are interquartile ranges. Shaded sectors indicate 95% confidence intervals. *P*-values in Fig 3B were corrected for *k* = 6 multiple comparisons using this formula *p*\* = 1 – (1 – *p*)*^k^*. Repeatability of individual behaviour within rounds was estimated with the intraclass correlation coefficient (ICC) in the *ICC* package in R (Wolak et al 2015).

## Results

In control experiments, we found that both tagged and untagged cockroaches preferred the shaded portion of the arena, showing no conspicuous differences in either tracking performance (Fig S1A) or speed (Fig S1B). The tagging treatment caused no significant differences in the distribution of aggregation index scores of groups of three animals (Fig S1C). Thus, the application of BEEtags did not appear to significantly alter naturalistic behaviour.

We measured the shade tracking performance of each of the six experimental cohorts in Round 1 six times each over successive days (Fig 1D). Cockroaches tracked the shaded sectors (Fig 1C, Movie S1), though they exhibited habituation to the stimulus over the course of 10 minutes (Fig S2). We chose a cut-off of 5 minutes for further trials to capture the highest performance shade tracking.

Cockroaches showed significant inter-individual variation in tracking performance (*p* < 10^−6^ by one-way ANOVA, df = 60, *F* = 3.599; Repeatability (estimated ICC [95% CI]) = 0.31 [0.20 – 0.44]; Fig 2A; Supplementary Table 1). The best tracking cockroaches avoided the light ~75% of the time, while the poorest trackers avoided it ~55% of the time. The distribution of tracking performance appeared to be roughly Gaussian. Individual shade tracking performance was stable across the 6 trials within Round 1, which spanned 6 days (Fig 2B). Notably, individual tracking performance, averaged across trials, was not correlated with the speed of individuals, averaged across trials (*r* = 0.13, *p* = 0.31; Fig S3). Because of this individual variation in tracking performance, cohorts also varied in their mean tracking performance across trials (Fig S4).

**Figure 2.**
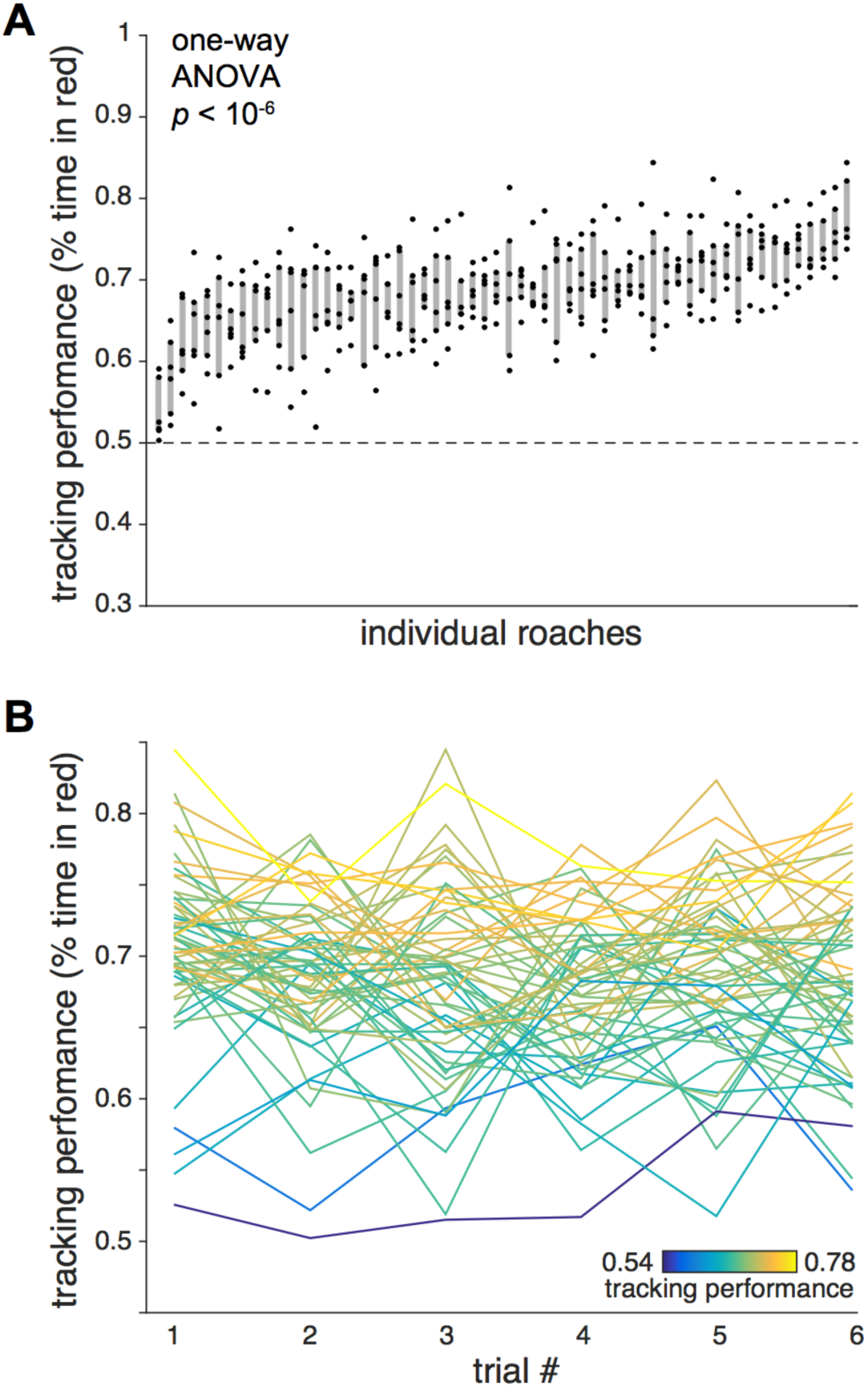
Cockroaches show stable idiosyncratic tracking performance during group behaviour. A) The percent of time that each cockroach spent in the shaded region (i.e. tracking performance) for each of the six Round 1 trials. A shaded region highlights the area between the first and third quartiles for each set of six data points. The cockroaches are sorted by average tracking performance. Dotted line shows the null expectation of a tracking performance of 0.5. **B)** Each cockroach’s tracking performance across all six trials of Round 1. Line colors indicate average tracking performance for that individual.

For Round 2, the cockroaches were placed in new experimental cohorts, based on their ranked tracking performance within their respective Round 1 cohorts (Fig 1D). The best performing individuals from each Round 1 cohort were placed together into a single Round 2 cohort, etc. As in Round 1, consistent interindividual variation in tracking was observed in Round 2 (*p* < 10^−6^ by one-way ANOVA, df = 58, *F* = 2.443; Repeatability = 0.20 [0.10 – 0.32]; Fig S5A; Supplementary Table 1), which persisted across days (Fig S5B). Likewise, cohorts in Round 2 varied in their average tracking performance (Fig S4B). Individual tracking performance in Round 2 was significantly correlated to individual tracking performance in Round 1 (*r* = 0.58, *p* < 0.0001; Fig 3A). Those individuals who tracked well in Round 1 continued to track well in Round 2 and those individuals who tracked poorly in Round 1 continued to track poorly in Round 2. The overall tracking performance of each cohort in Round 2 was not significantly different from a prediction based on the average performance of its members in Round 1 (df = 18, 2.37 > *t* > 0.062, 0.16 < *p* < 0.99 by multiple comparisons corrected t-test; Fig 3B). Thus, individual tracking performance in a group context appears to be robust to group composition.

**Figure 3.**
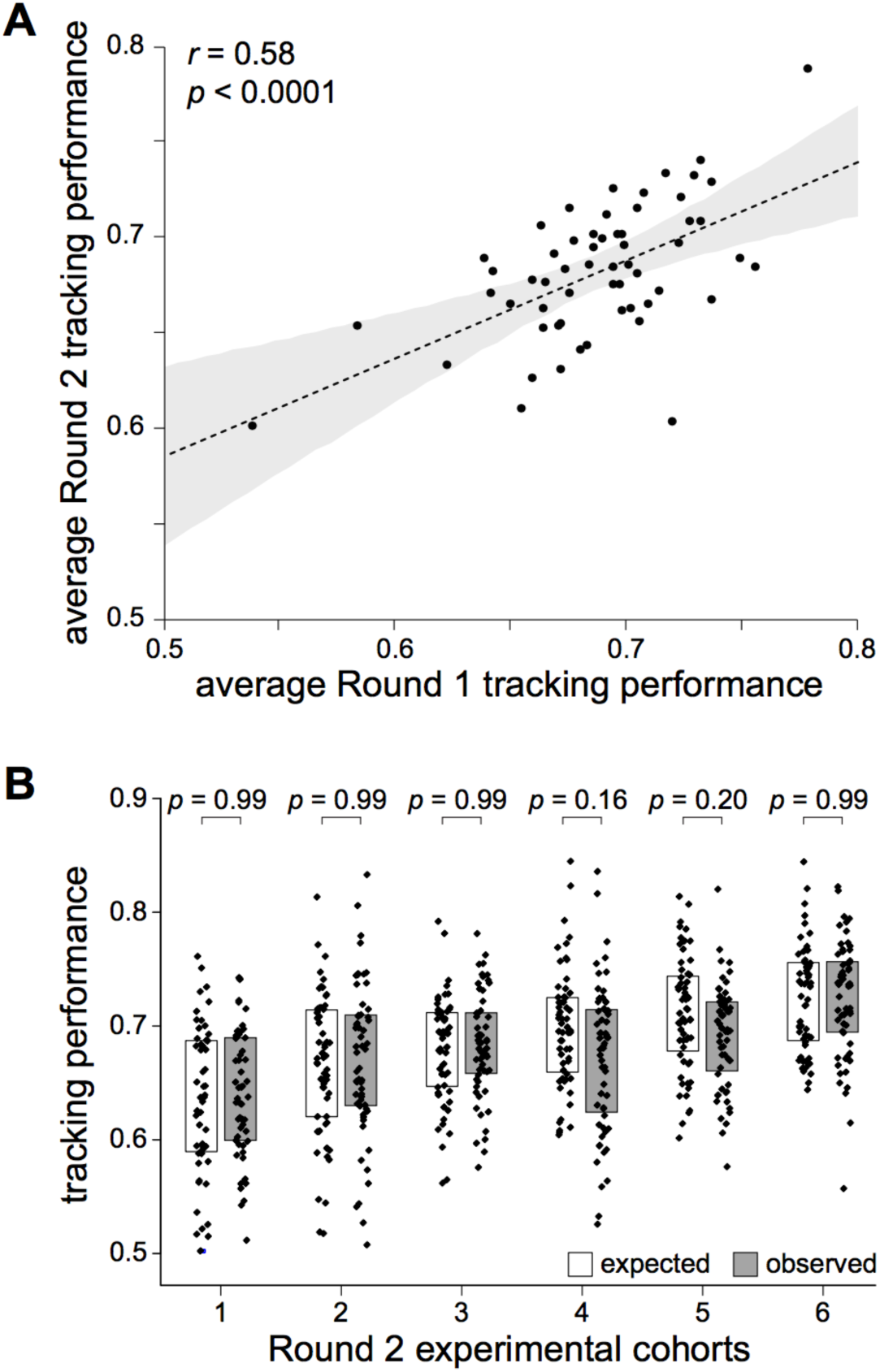
Idiosyncratic performance is robust to group composition. **A)** Scatter plot of average individual performance in Round 2 versus Round 1. Dotted line indicates linear fit. Shaded region indicates the 95% confidence interval on the linear regression. **B)** Expected (white) and observed (grey) distributions of individual trial tracking performance, by experimental cohort. Expected values are based on the per-trial performance of each cohort’s members in Round 1.

In Round 3, individuals were tested alone to see if the observed idiosyncratic behaviour, evident in groups, appears in a solitary context. All individuals were randomly assigned to six new housing groups of size ten (Fig 1D). From these housing cohorts, individuals were removed and tested alone under the same stimulus conditions as the earlier group tests (Movie S2). Concordant with previous results on collective light avoidance behaviour in cockroaches, the average tracking performance in solitary trials was significantly lower than in-group trials (Fig S6).

Cockroaches in Round 3 demonstrated consistent inter-individual variation in tracking performance in the solitary trials (*p* = 0.0015 by one-way ANOVA, df = 58, *F* = 1.821; Repeatability = 0.17 [0.05 - 0.32]; Fig 4A; Supplementary Table 1), which persisted over days (Fig S5C). Individual tracking performance (average of 4 Round 3 trials) in the solitary context was uncorrelated with tracking performance in the group context (average of 6 Round 2 trials) (*r* = 0.094, *p* = 0.48; Fig 4B). Thus the individual shade tracking performance observed in group contexts disappeared during solitary trials. In its place, new, consistent individual tracking performance levels appeared during solitary trials. As expected, the average tracking performance was lower in the solitary context than the group context (Fig 4A).

**Figure 4.**
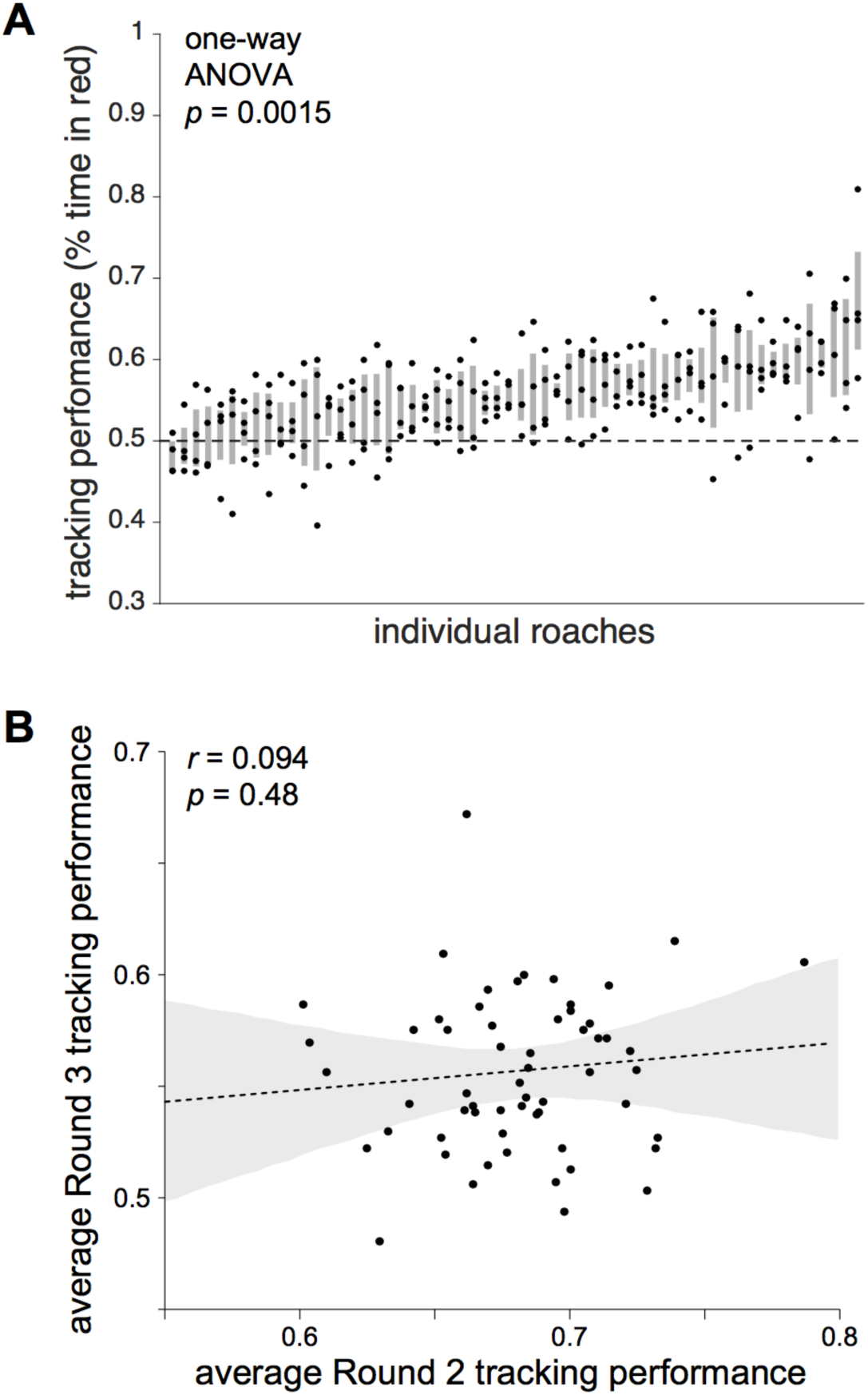
Solitary performance is not related to individual performance in groups. **A)** Tracking performance for each of the six Round 1 trials. Shaded regions show interquartile range. **B)** Scatter plot of average individual performance in Round 3 versus Round 2. Dotted line indicates linear fit. Shaded region indicates the 95% confidence interval on the linear regression.

The final experiments (Round 4) examined whether individual tracking performance levels would re-emerge when animals were restored to the group context during experiments. This was an important control when considering the possibility that over time and repeated manipulation the behaviour of the cockroaches may have drifted (Ridgel et al. 2003), which could trivially explain the lack of correlation between tracking performance between Round 3 and earlier Rounds. When individuals were randomly assigned to new experimental cohorts (Fig 1D), the observed individual tracking performances from Round 2 re-emerged. Average tracking performance was significantly correlated between Rounds 2 and 4 across individuals (*r* = 0.39, *p* = 0.0023; Fig 5A). Individual tracking performance in Round 4 was significantly correlated to Round 1 performance as well. Thus, all pairwise comparisons between Rounds of individual tracking performance in the group context were significantly correlated (Fig 5B). Conversely, the individual tracking performance in the solitary context was not significantly correlated with individual performance in any other Rounds (Fig 5B). As before, tracking performance shows significant inter-individual variation (*p* = 0.0032 by one-way ANOVA, df = 57, *F* = 1.834; Repeatability = 0.22 [0.06 – 0.40]; Fig S5D; Supplementary Table 1) and persistence across days (Fig S5E). As expected, cohorts in Round 4 differed in their average tracking performance (Fig S4C).

**Figure 5.**
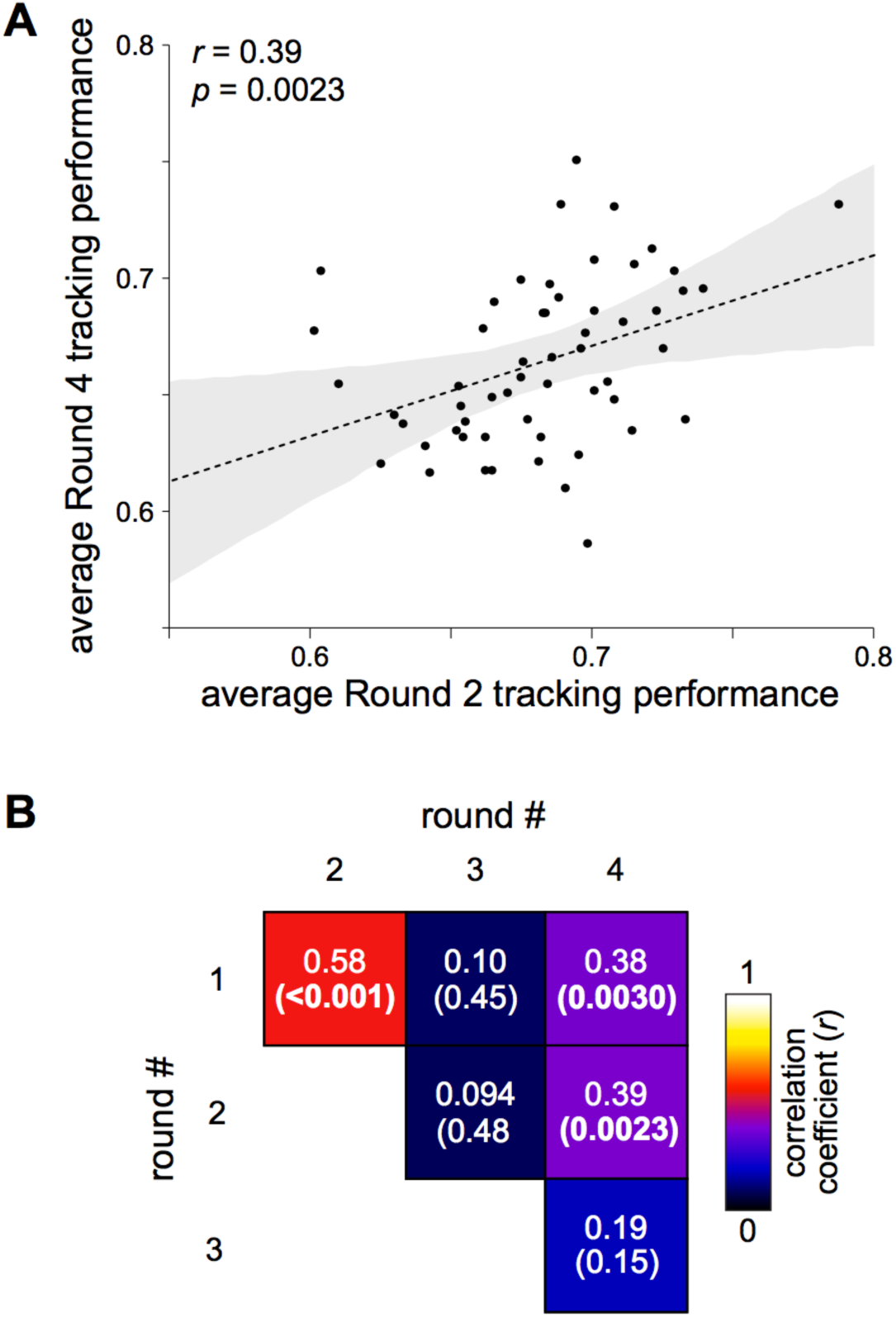
Returning individuals to groups restores group-specific tracking performance. **A)** Scatter plot of average individual tracking performance in Round 4 versus Round 2. Dotted line indicates linear fit. Shaded region indicates the 95% confidence interval on the linear regression. **B)** Pairwise Pearson correlation coefficients across individual cockroach tracking performances (averaged across trials), between all rounds. Numbers in each box show the correlation coefficient (above) and p-value (in parentheses below) for each pair of between-round correlations.

## Discussion

Our results demonstrate that cockroaches have individually consistent variation in shade tracking performance (Figs 1-2). We show that this idiosyncratic cockroach behaviour is robust to group composition (Fig 3) and is consistent over the course of several weeks (Fig 5), but surprisingly does not persist when cockroaches are tested in isolation from a group (Fig 4). Overall, these findings show that idiosyncratic behaviour is modulated by social context in cockroaches. While previous work has investigated how individual behavioural variation affects group performance in different classes of organisms (Millor et al., 2006; Canonge et al., 2009; Canonge et al., 2011; Burns et al., 2012; Briffa 2013; Marras et al., 2013; Pruitt & Keiser, 2014), it is less well understood how group membership influences individual behavioural performance.

These results have important implications for understanding the dynamics of collective decision making in animals. Despite increasing focus on both collective decision making (Arganda et al., 2012; Planas-Sitja et al., 2015) and consistent inter-individual variation (i.e. personality, Burns et al., 2012; Santos et al., 2014) in animals, the role of group heterogeneity in collective decision-making of animals, particularly in gregarious insects, remains a nascent research area. Where attempts have been made to understand the role of group heterogeneity in collective behaviour, this has typically been done by measuring personalities when individuals are separated (Briffa 2013; Brown & Irving 2013; Pruitt & Keiser 2014). This approach might not always be valid, however, because our results show that individual differences can be substantially modified by group context.

What drives individual behavioural variation in cockroach groups? One hypothesis for this variation could be the dichotomy between bold and shy (Frost et al., 2007; Sinn et al., 2008) or sitter-rover (de Belle & Sokolowski, 1987) personalities present in a wide variety of animals (Sih et al., 2004), since both higher activity level and higher portion of time in light could be considered characteristics of bold individuals. Especially for a mobile stimulus as used here, it is possible that more active individuals would perform better at tracking since they must actively search out the preferred stimulus. In our experiments, however, there was no relationship between activity level (i.e. velocity) and tracking performance (Fig S3, Movie S2).

Individual variation may also be produced dynamically in the presence of a social group, either by the presence of social hierarchies (Chase 1980), or by social niche differentiation (Bergmüller and Taborsky 2010). However, this does not appear to be the case in the experiments described here, since individuals did not shift behaviour in response to group composition (Fig 3) as would be expected from individual behavioural variation that emerges dynamically from the establishment of social hierarchies (Bell et al., 1979).

Although it is still possible that social niche differentiation plays a role in increasing behavioural variation among individual cockroaches, this effect would have to occur on a time scale of at least several weeks, since cockroaches housed together in new groups with lower behavioural variation (Round 2 above) for a week continuously showed no significant shift in their individual tracking performance. Alternatively, there may be a critical window for social niche construction, so that if an individual joins a niche sufficiently early in life, they will stay in that niche permanently even if re-grouped among individuals in the same niche. Even in this case, however, this social niche construction would seem to only apply in social contexts, since it disappeared when individuals were tested in isolation.

Another potential source of individual variation could be individual experience (Ravary et al 2007), for example arising from micro-environmental differences. However, for two reasons, we believe that environmental differences are unlikely to explain the sudden change in individual behaviours seen when animals are transferred to the solitary context. First we were careful to match their environmental circumstances during the experiments – matching their social conditions with constant group size housing, matching their visual experience by storing them in dark containers when not conducting experiments and making sure that all experimental handling was done by each experimenter across the whole cohort, rather than subsets of animals. Second, the re-emergence in Round 4 of the individual behaviours observed in Rounds 1 and 2 would be statistically improbable if environmental fluctuations explained the behavioural differences that occurred between Rounds 3 and 4.

As we observed, the correlation in individual shade tracking performance between solitary and group contexts is very weak, and could plausibly be less than zero (*r*=0.094 between rounds 2 and 3; 95% C.I.=[-0.19, 0.37] by bootstrap resampling). The lower correlation between solitary and group performance cannot be explained by sampling error alone, as all within-condition correlations (e.g. between pairs of Rounds 1, 2, 4) lie above the 95% C.I. of the solitary-group correlation.

A possible mechanism at play is that individuals vary in social cohesion (i.e. are more or less likely to stay next to other cockroaches), and this drives inter-individual variation in tracking performance, but only in the group context. Attraction to conspecifics plays an important role in many aspects of social behaviour (e.g. collective motion (Berdahl et al., 2013) or habitat selection (Stamps 1988)) and inter-individual variation in social cohesion can play an important role in structuring social behaviour (see, for example, Wey and Blumstein 2010). Since the group tracking performance is generally much higher than in separated individuals (Fig S6), individuals that are more likely to stay with a group of other cockroaches would likely have higher average tracking performance than individuals that ignore the presence of other cockroaches. However, the intensity of social cohesion among individual cockroaches would have no necessary bearing on performance when alone. This might explain the lack of correlation between individual performance levels in the solitary and group contexts (Fig 4). One way of testing this hypothesis in future work could be to examine inter-individual variation in levels of social cohesion, for example by measuring the amount of time cockroaches spend in proximity to a constrained group of other cockroaches in a behavioural arena. This test could be done in either the presence or absence of a light stimulus to investigate the interaction between social and visual stimuli.

This hypothesis highlights an important result of our experiments, namely that parameters driving individual behavioural performance in isolation may not have simple relationships to the parameters relevant for the same task when performed in a group. For example, in the group context, the probability of stopping next to another cockroach might be the single most important factor in determining tracking performance, while in isolation other behavioural parameters (e.g. velocity differences when in vs. out of the shade stimulus, etc.) may be much more relevant. If the correlation between individual performance in the group and solitary contexts is strictly zero, or negative, this implies that the cues driving shade tracking in both the solitary and group contexts (such as visual information) interact non-linearly with the cues present only in the group context (such as conspecific odour or tactile cues). If the interaction were linear, better-than-average exploitation of solitary cues would invariably be helpful in the group context, imparting a positive correlation. A slightly positive correlation could arise if significantly more linear weight were given to group-only cues; alternatively the presence of group-only cues could gate the processing of solitary cues. In humans, social context modulates numerous sensory channels including nociception (Krahé et al., 2013) and vision-touch integration (Heed et al., 2010).

In agreement with previous findings (Canonge et al. 2011, Salazar et al. 2013), we found that cockroach groups outperform individuals at tracking a shade stimulus (Fig S6). We hypothesize that this difference is due to the lack of information sharing typically associated with the presence of conspecifics and aggregation behaviour. The presence of conspecifics enhances public information sharing, which has been shown to be important for a variety of collective decision-making tasks (Miller et al., 2013), including locating shelters in cockroaches (Canonge et al., 2011).

One important consideration when interpreting the results of any longitudinal behaviour study in animals is the potential for time and physical injury to have influenced behaviour. Aging has been associated with neural degradation that affects not only gait mechanics but also the neural pathways associated with escape behaviours (Ridgel et al, 2003). While not statistically significant, our experiments showed a trend toward lower tracking performance in Round 4 when compared to Rounds 1 and 2 (Fig S6), as well as lower correlations between individual performance between Round 4 and the earlier Rounds 1 and 2 than was first observed between experimental Rounds 1 and 2 (Fig 5). These results suggest there may be at least weak levels of both behavioral drift and performance degradation.

Interestingly, cockroaches displayed the least individual consistency (i.e. the lowest repeatability) in the solitary context (Supplementary Table 1). This result is consistent with the emergence of more stable individual behavioural patterns within a group context, an affect that has been observed in solitary ant queens (Fewell and Page 1999), and has been suggested to potentially play an important role in the evolution of division of labour.

Most broadly, our results highlight the importance of considering group context when examining behaviour in animals. For animals exhibiting any degree of social behaviour, from occasionally gregarious animals to the highly eusocial insects, groups are not only composed of individuals of different types, but may also play an active role in modulating and creating individual variation in collective behaviours.

## Author Contributions

JDC, ADS, DA, SDH and BLdB designed the experiments. JDC, ADS, DA and SDH carried out the experiments. JDC, ADS, DA, SDH and BLdB analyzed the paper and wrote the manuscript. SEC and WMC designed preliminary experiments and analyzed preliminary data. MWB and BLdB supervised preliminary experiments.

## Acknowledgements

We thank Clark Magnan, Ed Soucy and Joel Greenwood for help with implementing experiments.

## Supplementary materials

**Movie S1** – *Example group experiment* – Shade-tracking behaviour of a group of nine cockroaches. Movie was taken after the end of data collection as an example. Recorded with a digital camera with a rolling shutter RGB sensor.

https://youtu.be/SYe5h4QDDE4

**Movie S2** – *Example solitary experiment* – Shade-tracking behaviour of a single cockroach from the solitary experiments (Round 3). Recorded with a digital camera with a rolling shutter RGB sensor.

https://youtu.be/hw-zffKJH4s

**Movie S3** – *Automated tracking follows individual paths over time in a dynamic stimulus* – Individual tracked (green) and interpolated (red) positions of ten cockroaches over time. In the second half of the movie, individual colors show traces of individual position for ten frames before the current one. Movie was captured at 7 frames per second and is played back at 30 frames per second.

https://youtu.be/-M4Rtk1cEVg

**Figure S1.**
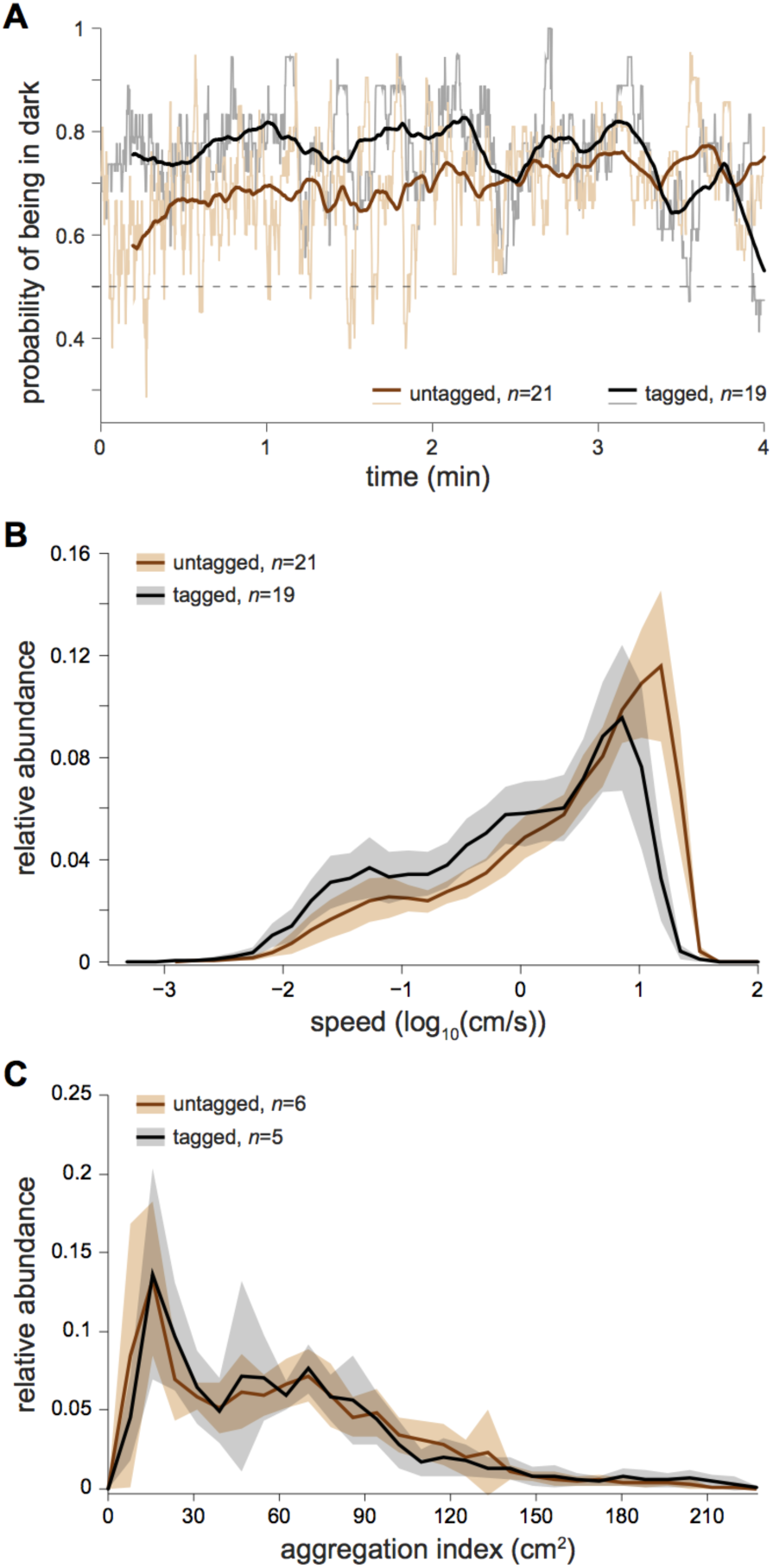
Tagging does not disrupt cockroach behaviour. **A)** Tracking performance, over time, **B)** Normalized histograms of speed (*p* = 0.67 by Kolmogorov-Smirnov test), and **C)** Normalized histograms of the aggregation index (*p* = 0.76 by Kolmogorov-Smirnov test) for tagged (black) and untagged (brown) cockroaches.

**Figure S2.**
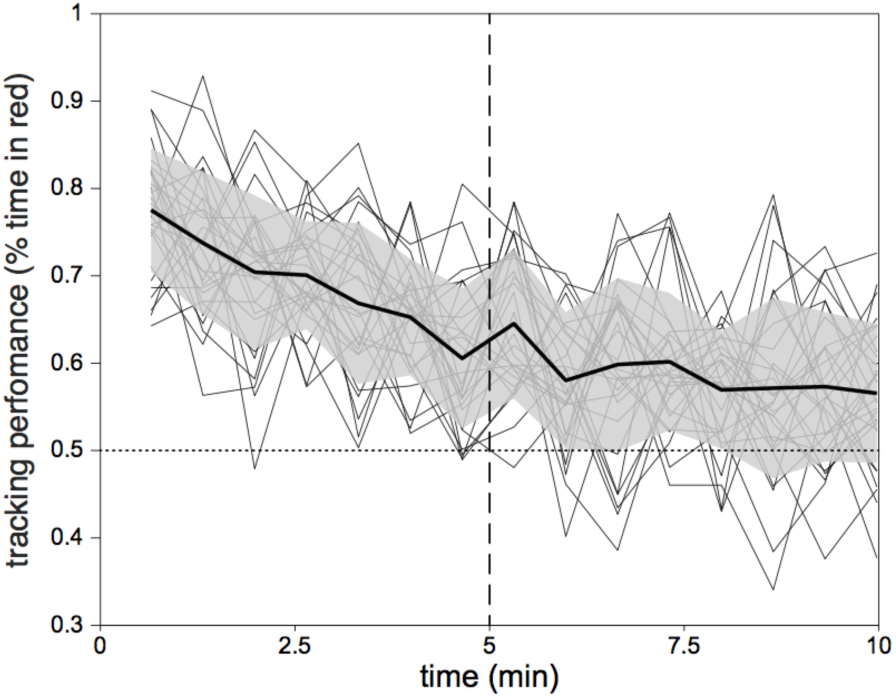
Cockroaches habituated to the light stimulus. A plot of group tracking performance vs time across trials in Round 1. Thin black lines show group tracking performance from individual trials. Thick line shows the average group tracking performance vs time across trials, and grey shaded regions indicate the mean +/− one standard deviation.

**Figure S3.**
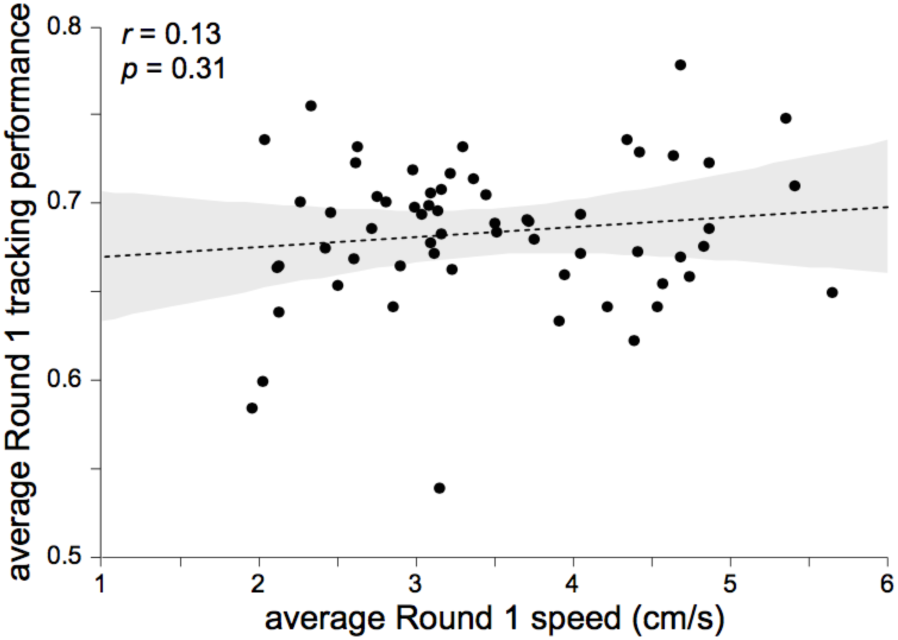
More active cockroaches are not better trackers. Individually-averaged tracking performance vs. speed in Round 1.

**Figure S4.**
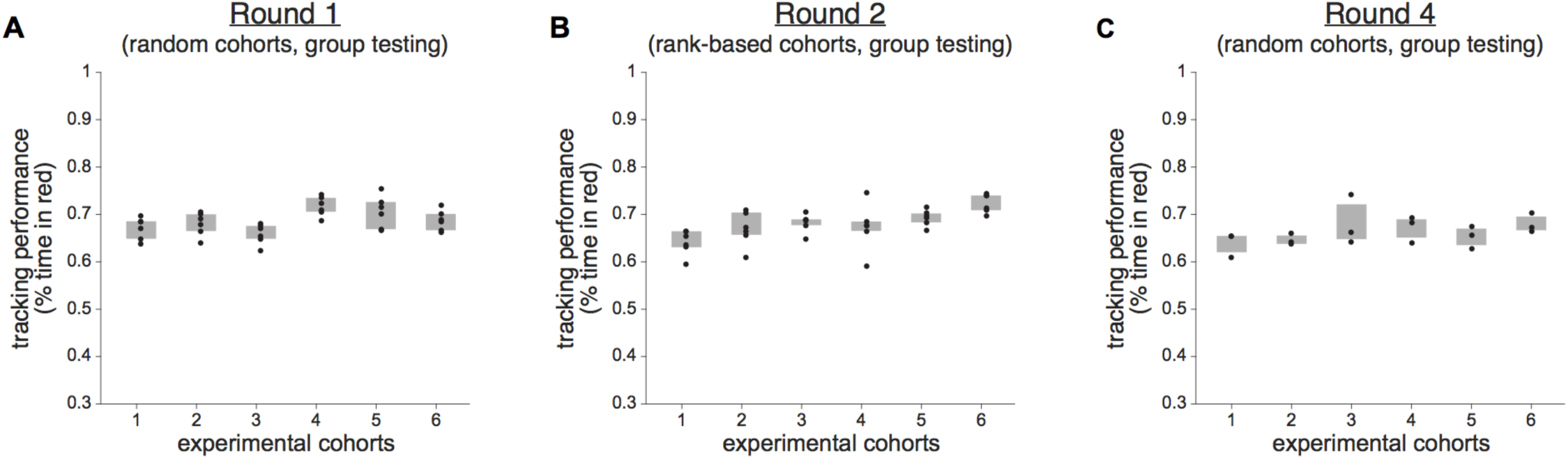
Cohorts vary in average group tracking performance. Group tracking performance by cohort in **A)** Round 1, **B)** Round 2, and **C)** Round 4. Black dots represent average tracking performance from individual trials and shaded regions indicate interquartile range.

**Figure S5.**
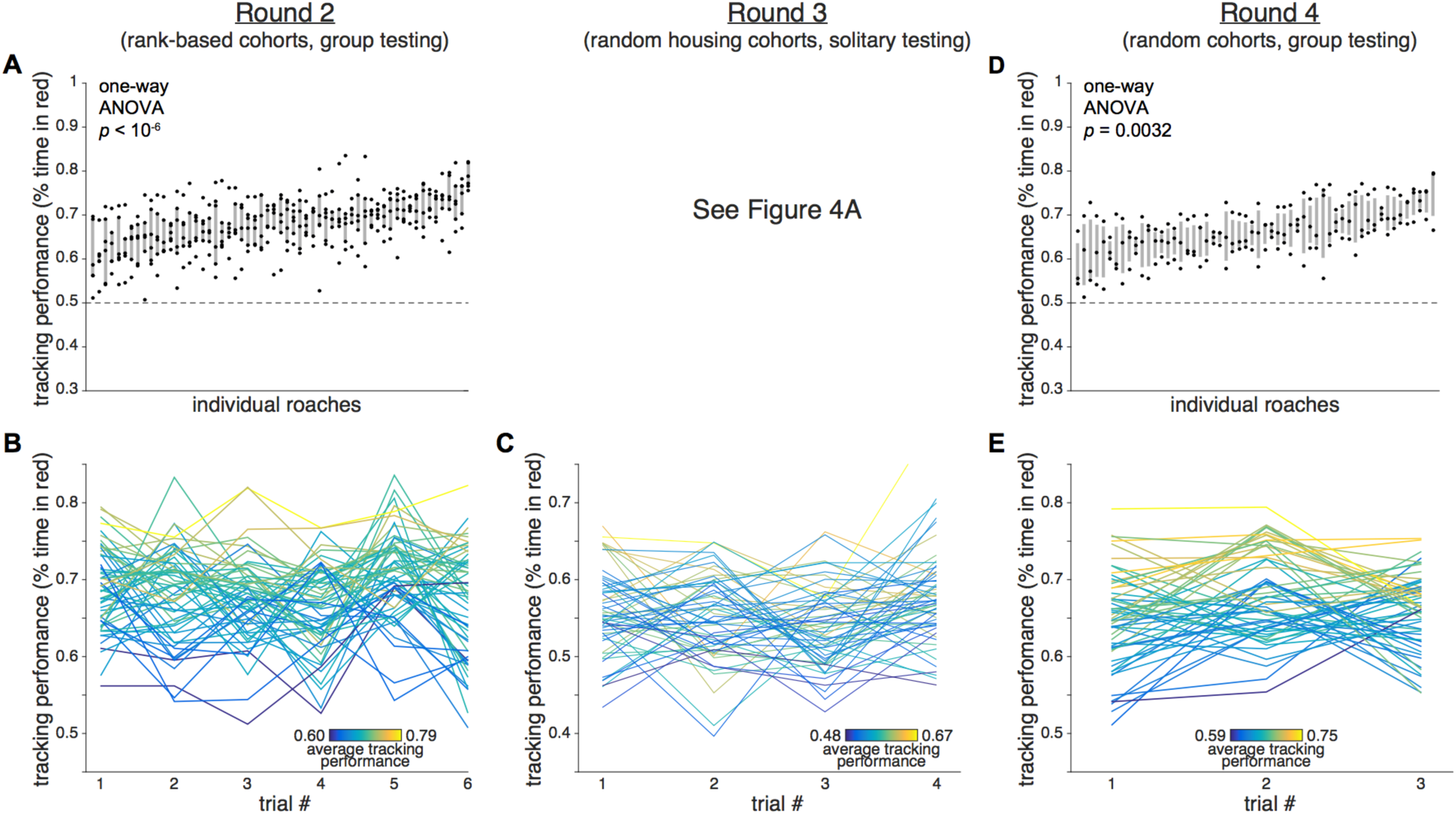
Cockroaches display consistent idiosyncrasy in tracking performance in all experimental conditions. **A)** Tracking performance for each of the six Round 2 trials. Shaded regions show interquartile range. **B)** Each cockroach’s tracking performance across all six trials of Round 2. **C)** Each cockroach’s tracking performance across all six trials of Round 3. **D)** Tracking performance for each of the three Round 4 trials. Shaded regions show interquartile range. **E)** Each cockroach’s tracking performance across all three trials of Round 4. Line colors in **B**, **C**, and **E** indicate average tracking performance for that individual.

**Figure S6.**
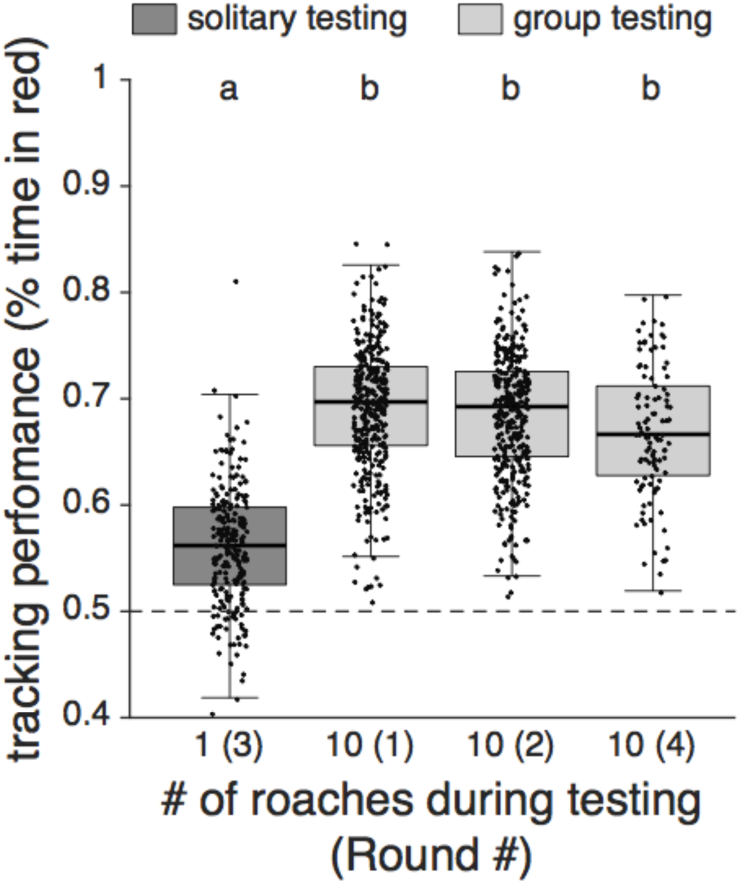
Cockroach groups track better than individuals. Tracking performance by experimental round. Black dots show tracking performance from a single cockroach in a single trial, grey region shows the interquartile range. Letters indicate statistically significant groups by pairwise t-tests.

